# For colonization success, should hosts and microbes travel alone, together, or swap partners along the way?

**DOI:** 10.1101/2025.10.27.684839

**Authors:** Takuji Usui, Jingcheng Yu, Megan E. Frederickson

**Author notes:** Corresponding author: Takuji Usui (Department of Ecology & Evolutionary Biology, University of Toronto, 25 Willcocks Street, Toronto, ON, M5S 3B2, Canada. Co-First Authors (Takuji Usui, Jingcheng Yu).

## Abstract

Understanding what determines colonization success is a key challenge of global change ecology. Microbiomes that enhance host performance are likely to be co-introduced with hosts during colonization due to their intimate association. Yet, it is unclear how co-introduced microbes will impact host colonization, as both the microbiome and its effects could vary upon introduction into a new habitat. Here, we used duckweeds —a cosmopolitan, freshwater angiosperm— and their microbiome to track colonization dynamics during an experimental co-introduction in the wild. Notably, we found that host performance was substantially reduced when plants were co-introduced with microbes non-local to the introduced habitat, relative to hosts with a local microbiome. Moreover, negative impacts from the non-local microbiome persisted for multiple host generations despite a rapid turnover in microbiome composition during colonization. Overall, we suggest that considering the co-introduced microbiome can help predict host colonization success under global change.

## INTRODUCTION

The movement of species due to both biological invasions and climate change-induced range shifts is a hallmark of global change (Chen et al. 2011; Iseli et al. 2023). As such, a critical goal of contemporary ecology is to understand what makes certain organisms more successful at colonizing and establishing in new habitats (Gioria et al. 2023). While the importance of antagonistic interactions (e.g., competition) in shaping colonization and establishment has long been recognized (Cadotte 2007; Case 1990), there is growing recognition that mutualistic interactions also determine colonization success (Fowler et al. 2023; Moles et al. 2022; Shaw et al. 2021). Most macroscopic organisms host a microbiome that frequently engages in reciprocally beneficial interactions (i.e., mutualisms) (Frederickson 2013; Rodriguez et al. 2009). Because the microbiome is intimately associated with a host, with microbes living on or in host tissues, these mutualistic interactions between hosts and microbes could be particularly important as both hosts and their microbiome may frequently be co-introduced to new environments (Le Roux et al. 2017; Mestre et al. 2024). Yet, much remains unclear about the role and importance of co-introduced microbes for host performance during colonization.

On one hand, mutualistic microbes that are co-introduced with hosts could provide immediate benefits that increase host performance in the new habitat, for example, through improved stress tolerance or protection against pathogens or other natural enemies (Afkhami et al. 2014; Benning & Moeller 2021). Furthermore, beneficial interactions with microbes can not only increase mean population growth but can also stabilize demographic variance in host populations (Fowler et al. 2024). As the early establishment phase of colonization–where population sizes are small and vulnerable to local extinction–represents a key bottleneck for introduced populations (Catford et al. 2016; Van Kleunen et al. 2018), initial communities of co-introduced microbes could therefore have a large influence on host persistence. In the well-studied legume-rhizobia mutualism, microbial sequencing data suggest that invasive legumes are often co-introduced with their nitrogen-fixing bacterial symbionts (Ndlovu et al. 2013; Rodríguez-Echeverría 2010; Warrington et al. 2019), potentially improving host colonization success. Such co-introductions may be especially important for hosts engaging in more specialized mutualisms, such as the legume-rhizobium mutualisms, in which a lack of appropriate partners in the introduced range may inhibit host establishment and spread (Harrison et al. 2018; Nathan et al. 2023; Simonsen et al. 2017).

On the other hand, if microbial partners are locally adapted to their environments (i.e., a microbe genotype by environment, or G_microbe_ x E, interaction), microbes that are local to the introduced habitat may confer greater host benefits relative to co-introduced, non-local microbes (Hui & Richardson 2017). Additionally, as host-microbiome interactions are often context-dependent (Chamberlain et al. 2014), the magnitude of host benefits derived from co-introduced microbes may also change upon introduction into a new habitat (Le Roux et al. 2017; Traveset & Richardson 2014). For example, if the interaction between the host and microbiome depends on the environment (i.e., a G_host_ x G_microbe_ x E interaction), mismatches between co-introduced hosts and microbes to their new environment could reduce host benefits, and in severe cases, may shift the host-microbiome relationship across the mutualism-parasitism continuum (Drew et al. 2021; Hoeksema & Bruna 2015). In such cases, generalist hosts might benefit from switching partners to recruit local microbes that are present in the introduced habitat (i.e., ‘ecological fitting’ *sensu* Le Roux et al. 2017).

Yet, local microbial communities may still lower the performance of introduced hosts, for example, through the presence of novel pathogens or a lower abundance of beneficial partners in the new habitat (Lankau & Keymer 2016; Nuñez et al. 2009). Host-microbiome coevolution could also cause the recruitment and effects of microbes to depend on the host’s genetic background (i.e., a G_host_ x G_microbe_ interaction) (Brown et al. 2020; Nuismer & Gandon 2008; Wagner et al. 2016), such that non-local host genotypes may associate with different microbes or fail to derive the same benefits as local host genotypes. Additionally, the recruitment of local microbes may be further complicated if hosts travel with their co-introduced microbiome. For example, past research has found that microbes that are initially present on hosts can modify resource availability or the host environment (Debray et al. 2022), thereby inhibiting or altering the assembly of subsequent microbial symbionts through competitive exclusion or priority effects (Boyle et al. 2021; Chappell et al. 2022). This means that the initial co-introduction of hosts with their microbiome could also have cascading effects on the recruitment —and therefore influence— of local microbes during colonization.

Few experiments manipulate the co-introduction of hosts and microbiomes and test their effects on host performance in the wild, so it remains an open question whether hosts should travel alone, together with their microbiome, or swap partners along the way for successful colonization. Here, we use the common duckweed (*Lemna japonica*) and its microbiome as a model community for testing how microbes alter host performance during an experimental introduction into a common pond (Fig. 1). By manipulating the introduced host genotype (G_host_) and the identity of the initial, co-introduced microbiome (G_microbe_), we tested if: (1) host performance differed when hosts were co-introduced with or without local and non-local microbes; (2) if the presence of co-introduced microbes altered host recruitment and association with local microbes; and (3) if host genetic background altered the recruitment and association of microbes during colonization.

**FIGURE 1.**
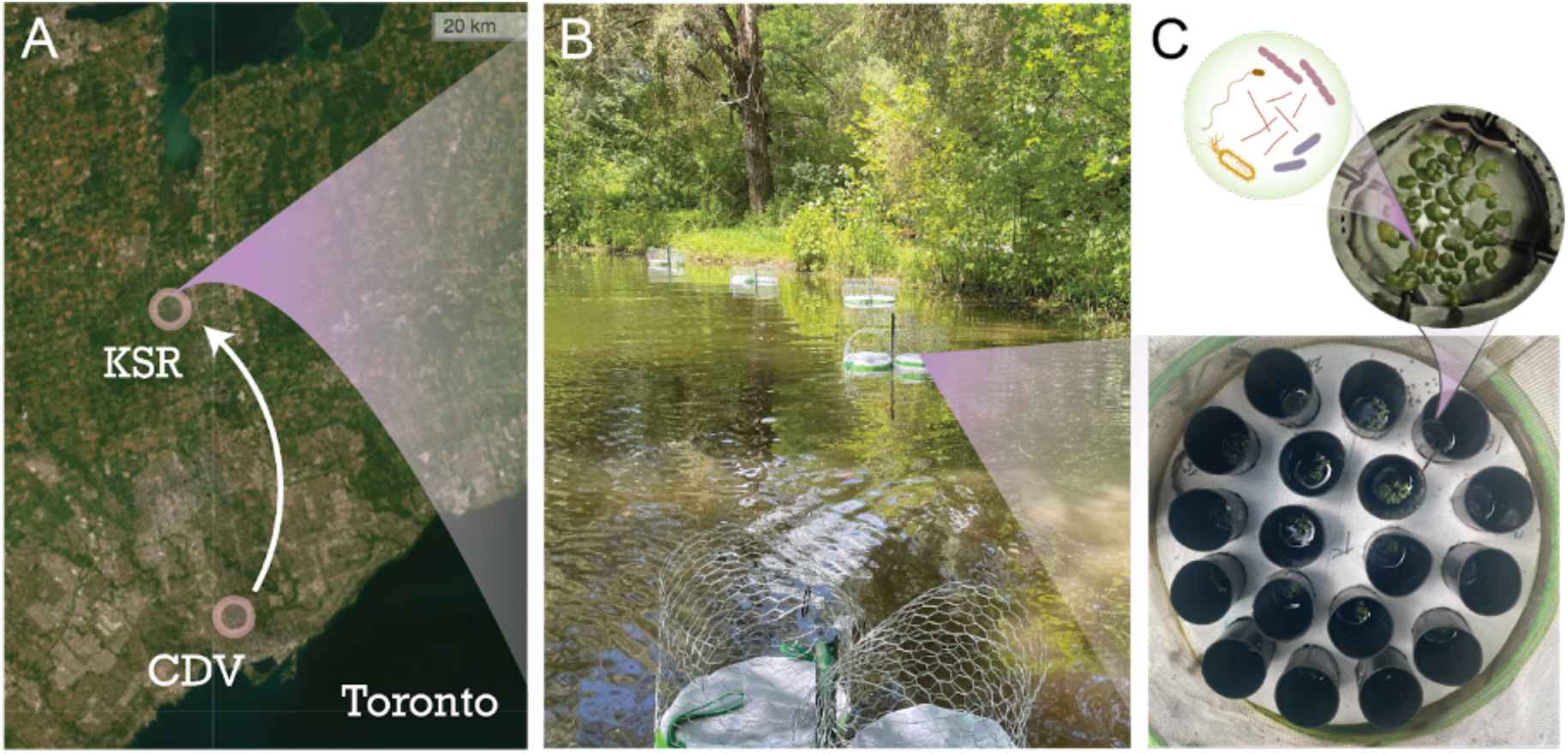
Experimental co-introduction of duckweeds and their microbiome in a common pond. (A) Sampling sites of duckweeds and microbes at the Koffler Scientific Reserve (KSR; –79.518, 44.035) and Cedarvale Park (CDV; –79.419, 43.690), ON, Canada. (B) Platforms anchored around the perimeter of a single pond at KSR (C) Each platform (*r* = 18.5 cm) housed 18 replicate cone-tainers (*r* = 2.3 cm) in which we seeded *L. japonica* plants with spatially randomized microbial treatment groups within each platform.

## MATERIALS & METHODS

### Collection and manipulation of duckweeds and microbiome

Duckweeds are floating, freshwater macrophytes that host a generalist and often mutualistic microbiome that resembles the microbiome of terrestrial angiosperms (Acosta et al. 2020; O’Brien et al. 2020; Tan et al. 2021). Considered one of the fastest growing angiosperms (Laird & Barks 2018), duckweeds can tolerate a wide range of environments and are frequently dispersed between ponds by waterfowl (Coughlan et al. 2015, 2017). These traits contribute to their widespread distribution, with several species of *Lemna* considered invasive (Andrade-Pereira 2024). *Lemna japonica* is a recently identified cryptic hybrid of *Lemna minor* and *Lemna turionifera* (Braglia et al. 2021), although its geographical distribution is currently unknown. We used two host *L. japonica* genotypes sampled from two geographically distinct ponds in the Greater Toronto Area, Ontario, between 2017 and 2021: Koffler Scientific Reserve (KSR; –79.518, 44.035) and Cedarvale Park (CDV; –79.419, 43.690) (Fig. 1A). As duckweeds grow clonally and have one of the lowest per base pair mutation rates among multicellular eukaryotes (Sandler et al. 2020), there is very little segregating genetic diversity within sites (i.e., ponds) and most genetic differentiation occurs among sites (Cole & Voskuil 1996; Ho et al. 2019; Schmid et al. 2024). Our KSR and CDV duckweeds are distinct clonal lineages based on long-read SNP data (Usui, O’Brien, Harkess, Wright, & Frederickson, unpublished data) and we therefore refer to them as distinct host genotypes (Laurich et al. 2024; O’Brien et al. 2020).

We first sterilized plants of each genotype following a standardized protocol described in Laurich et al. (2019). Briefly, we vortexed sampled plants in phosphate-buffered saline, immersed plants in 1% sodium hypochlorite, and then maintained sterile plants in 200mL of autoclaved, Krazčič growth media (Krazčič et al. 1995) inside a growth chamber (set at a constant temperature of 22 ℃ with a 16:8 hour light:dark cycle). Plants were periodically assessed for sterility by enriching the growth media with 5g/L sucrose and 1g/L yeast extract to check for microbial growth. If any microbial growth was observed, plants were re-sterilized following the same procedure. We also collected the natural, host-associated microbiome of our KSR and CDV duckweeds by sampling around 100 duckweed plants and pond water from the same two sites in July, 2024. To isolate the microbiome, sampled plants and pond water were vigorously shaken and then filtered through 2µm filter paper to remove plants and large debris. We then centrifuged the filtered sample into a pellet (3000 rcf for 10 minutes) and resuspended the microbial inocula with 200mL of autoclaved, Krazčič growth media (Laurich et al. 2024; Wei & Tan 2023).

Before the pond experiment, we split each of the two host genotypes into three initial microbial treatment groups, with each host genotype grown: (1) in sterile condition in fresh, autoclaved growth media (i.e., ‘uninoculated’); (2) in growth media inoculated with microbes collected from its home site; or (3) in growth media inoculated with microbes collected from the alternate site (i.e., a fully factorial G_host_ x G_microbe_ manipulation). Plants were cultured under these microbial treatments for 14 days to ensure that the microbial community had fully interacted with duckweed hosts (Acosta et al. 2020). Before using these plants as the source population for experimental introductions, we re-checked the sterility of uninoculated plants by loading 20µL of growth media onto plates containing LB agar medium. After 4 days, we then visually confirmed that there was no microbial growth on these plates.

### Experimental co-introductions and host data collection

We conducted a common pond experiment in which we tracked the colonization dynamics of introduced hosts and their microbiome in a natural pond at one of the original sampling sites at the Koffler Scientific Reserve (KSR; between July and August, 2024) (Fig. 1B). We therefore refer to host genotypes and microbes originating from KSR as ‘home’ (i.e., local), and host genotypes and microbes originating from Cedarvale (CDV) as ‘away’ (i.e., non-local) hereafter. While both sites are located within the range of *L. japonica*, the co-introduction of away plants and microbes into the common pond serves to simulate the natural dispersal and colonization of duckweeds that occurs between distinct ponds via waterfowl (Coughlan et al. 2015, 2017).

For the experiment, we deployed 12 floating platforms made of polystyrene following Subramanian & Turcotte (2020) (Fig. 1C). Each platform held 18 replicate cone-tainers (*r* = 2.3 cm; Ray Leach Cone-Tainers, OR, USA) with holes drilled on the underside to allow for water exchange. Platforms were enclosed in a butterfly net surrounded by galvanized fencing for protection. We secured two platforms each to six fence posts that were anchored to a fixed position in the pond substrate. Fence posts were spaced at least 8m apart from each other along the edge of the pond where duckweeds typically reside (Fig. 1B). The platforms were deployed in the pond for 2 weeks before the introduction of host plants to ensure adequate water exchange and mixing. We then seeded 8-10 duckweed individuals per cone-tainer on day 0 of the experiment. Within each platform, host-microbe combinations were spatially randomized (*N* = 36 replicate populations per host-microbe combination for a total of *N* = 216 populations across all treatment groups). During the experiment, we tracked host performance (i.e., growth rate) by censusing all populations on days 2, 4, 7, 9, 11, and 14. Population censuses were disturbed for *N* = 14 replicates in one platform (6% of all replicates) which was destroyed by a sole green frog (*Lithobates clamitans*). These replicates were removed from all subsequent data collection and analyses.

### Collection and sequencing of the microbiome

To characterize changes in the duckweed microbiome during colonization, we sampled host plants: (1) from the source population in the lab prior to pond introduction on day 0; and (2) from a subset of populations on days 4, 9, and 14 from the common pond. At each time point, we destructively sampled and froze all plants from 3-6 replicate populations per host-microbe combination. We extracted microbial DNA from frozen plant tissue using the PureLink Genomic DNA Mini Kit (Thermo Fisher Scientific, Waltham, MA, USA). We then sent >10ng of DNA per sample to Génome Québec (McGill University, QC, Canada) for PCR amplification of the 16s rRNA gene (V3-V4 region; 341F/805R primers; Table S1) followed by paired-end, 250 base pair sequencing on an Illumina MiSeq System (San Diego, CA, USA). We received demultiplexed reads with an average of 144,100 reads per sample.

We then used the QIIME2 software (v2022.2) (Bolyen et al. 2019) to process reads. We first trimmed adapters and primers with *cutadapt* (Martin 2011) and then filtered for sequencing errors and assigned amplicon sequence variants (ASVs) to paired reads with *DADA2* (Callahan et al. 2016). We assigned taxonomic classifications using the naive Bayesian *classify-sklearn* method pre-trained on the Greengenes2 database (McDonald et al. 2024), limiting identifications to a confidence threshold of ≥70%, and filtering out plant chloroplast and mitochondrial sequences. Finally, we removed very rare ASVs (<10 reads across all samples) and rarefied the dataset to the number of reads in the smallest sample (*N* = 31,403 reads) after visualization of a rarefaction curve. We obtained a final dataset of 114 samples with 2765 unique ASVs across 353 bacterial families.

### Statistical analyses

All linear-mixed effects models were conducted in a Bayesian framework using *MCMCglmm* (Hadfield 2010) in R (v4.4.3; R Development Core Team, 2025). For all models, we used a weakly informative, inverse Wishart prior with *V* = 1 and *nu* = 0.002 for random effects (Feller & Gelman 2015), running models for 50,000 interactions, discarding the first 1,000 iterations as burn-in, and sampling every 10th iteration. We report posterior means, 95% credible intervals (i.e., the smallest interval which captures 95% of the posterior values), and the *p*MCMC (i.e., an analogue of the frequentist two-tailed *p*-value).

To test for differences in host performance during colonization, we modelled the number of individuals as the response variable, and a three-way interaction between time (days), host genotype (home or away genotypes), and initial microbial treatment (uninoculated, home (i.e., KSR) microbes, or away (i.e., CDV) microbes) as fixed predictor variables. We removed data from populations in which censuses were cut short by the destructive harvesting of the plant microbiome (although see Fig. S1 and Table S5 for the full dataset), and included nested random effects of paired platform ID, platform ID, and cone-tainer ID.

For the microbiome, we first tested for differences among the initial microbiome of lab-cultured host populations on day 0 prior to introduction. To do so, we estimated weighted UniFrac distances between each pair of samples using *rbiom* (Smith 2025), which measures the pairwise phylogenetic distance between communities while accounting for the relative abundance of each taxon (Lozupone et al. 2011). We conducted a PERMANOVA on these weighted UniFrac distances, fitting an interaction between host genotype and microbial treatment as fixed predictors using the *adonis2* function in *vegan* (Okansen et al. 2025).

Additionally, we explored how representative these lab-cultured plant microbiomes were of natural, field-sampled plant microbiomes (*N* = 6) from the home site on day 0. Specifically, we tested if microbial diversity differed between lab-cultured and field-sampled hosts by fitting Shannon’s diversity (H’) as the response variable, and sample type (lab or field) as the fixed predictor in a linear model. We also tested if home plants inoculated with home microbes were most similar in community composition to home field samples, by fitting weighted UniFrac distances (between lab and field samples) as the response variable, and an interaction between host genotype and initial microbial treatment as the fixed predictor.

To test for microbiome changes during colonization, we calculated weighted UniFrac distances between all samples from the common pond on days 4, 9, and 14, and conducted a PERMANOVA with a three-way interaction between host genotype, initial microbial treatment, and time as fixed predictors. We then tested if host microbiomes became more or less similar to each other during colonization, by fitting weighted UniFrac distances as the response and time as a fixed predictor in a linear model. Upon finding treatment differences in host performance, we also conducted post-hoc PERMANOVAs on data subset to uninoculated or inoculated hosts. Specifically, we tested: (1) if microbes that uninoculated hosts acquired during colonization depended on host genotype (i.e., interaction between host genotype and time as a fixed predictor); and (2) if microbes that inoculated hosts acquired during colonization depended on the initial microbiome (i.e., interaction between initial microbial treatment and time as a fixed predictor).

Finally, we tested if variation in the microbiome during colonization contributed to differences in host performance. We first tested if the first and second PCoA axes –which captured 60.7% and 15.3% of the variation in the microbiome of hosts after introduction, respectively– explained differences in host performance. Specifically, we fit host growth rate as the response variable and fit the first or second PCoA axes as a fixed predictor in separate linear mixed-effects models, with paired platform ID, platform ID, and sampling day as random effects. As PCoA values were available for every plant population that was destructively harvested on days 4, 9, and 14, we estimated host growth rates for each population up until the day of harvest. We found a single, extreme outlier for host growth rate on day 4 and thus we removed this sample prior to analyses. The first PCoA axis was strongly right-skewed and so was transformed before analysis by adding a minimum constant value to make all values positive and then applying a log-transformation (i.e., a log-shift transformation). To identify specific microbial taxa associated with host performance, we additionally ran a phylogenetic factorization on these microbiomes using *Phylofactor* (Washburne *et al*. 2017). This approach incorporates both the phylogenetic and compositional structure of microbiomes (using isometric log-ratio (ILR) transforms of abundance) to iteratively partition a phylogeny at varying taxonomic depths, identifying clades that are most important in its association with a given variable (i.e., host performance) (Washburne et al. 2017). Thus, one advantage to traditional differential abundance tests is that the taxonomic scale of investigation is a biologically meaningful output of the model rather than a parameter that is arbitrarily chosen prior to analysis.

## RESULTS

### Co-introduced microbiomes alter the trajectory of host growth

We found that host performance varied with both the initial microbial treatment and host genotype (Fig. 2). First, among host plants that were initially inoculated with a microbiome, hosts introduced with home (KSR) microbes generally grew faster than those introduced with away (CDV) microbes, regardless of host genotype (Fig. 2B). Notably, compared to home plants with home microbes, hosts introduced with away microbes had a reduced growth rate for both home (mean reduction = –0.144, –0.238 to –0.044 N/day, *p*MCMC = 0.002; Table S2) and away host genotypes (mean reduction = –0.178, –0.273 to –0.081 N/day, *p*MCMC <0.001; Table S3). Second, among hosts that were initially microbe-free, we found that the growth trajectory of introduced populations depended on host genotype (Fig. 2B). We found that home plants with initially no microbiome had the highest estimated growth rate (0.806, 0.736 to 0.873 N/day; Fig. 2B), while in comparison, the growth of initially microbe-free away plants was significantly lower (mean reduction = –0.260, –0.356 to –0.161 N/day, *p*MCMC < 0.001; Table S4). In fact, away plants introduced without a microbiome were estimated to have the slowest growth rate out of any treatment group (0.546, 0.380 to 0.712 N/day; Fig. 2B). Taken together, host performance is therefore highest for hosts introduced with home microbes and for home plants that are initially microbe-free, while host performance is lowest for hosts introduced with away microbes and for away plants that are initially microbe-free. We found that these results remained consistent in models that included growth data obtained from populations harvested during the experiment (Fig. S1 and Table S5).

**FIGURE 2.**
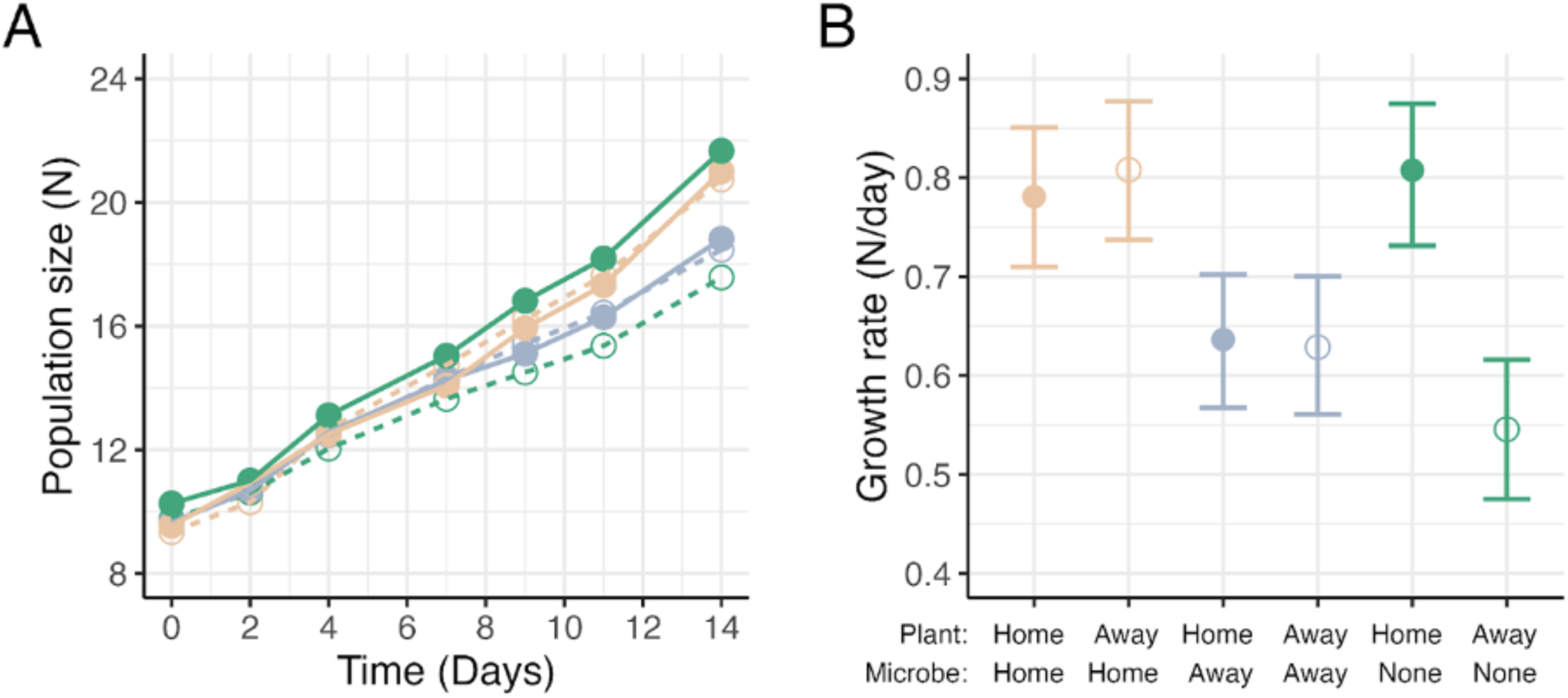
Host performance differs with host plant genotype (home = filled circles and solid lines; away = open circles and dashed lines) and initial microbial treatment (home = orange; away = blue; uninoculated = green). (A) Lines represent the growth trajectory of introduced host populations over time where circles represent the mean population size at a given day for each host genotype and microbial treatment group. (B) Mean population growth rate (N/day) and 95% CIs for each host and microbe combination, as estimated from linear mixed-effects models.

### Initial microbiomes differ mostly by host genotype

Prior to pond introduction, we found that the plant microbiome comprised a total of 876 ASVs from 105 bacterial families. Of these, 136 (15.5%) ASVs were unique to plants cultured with local (KSR) microbes, while 141 (16.1%) ASVs were unique to away (CDV) microbes (12.4% and 5.7% at the family-level, respectively; Fig. 3A; Fig. S2). While we found distinct ASVs between the home and away microbiome, ordination analyses on weighted UniFrac distances did not show a significant effect of microbial treatment (PERMANOVA: F_(1,8)_ = 1.283, R^2^ = 0.098, *p* = 0.107; Table S6). Instead, the microbiome differed significantly by host genotype (PERMANOVA: F_(1,8)_ = 7.904, R^2^ = 0.402, *p* < 0.001; Fig. 3B). We also did not detect a significant microbe by host genotype interaction (PERMANOVA: F_(1,8)_ = 1.831, R^2^ = 0.093, *p* = 0.121; Table S6), although this may reflect limited statistical power due to a relatively small sample size (N = 12) of initial samples.

**FIGURE 3.**
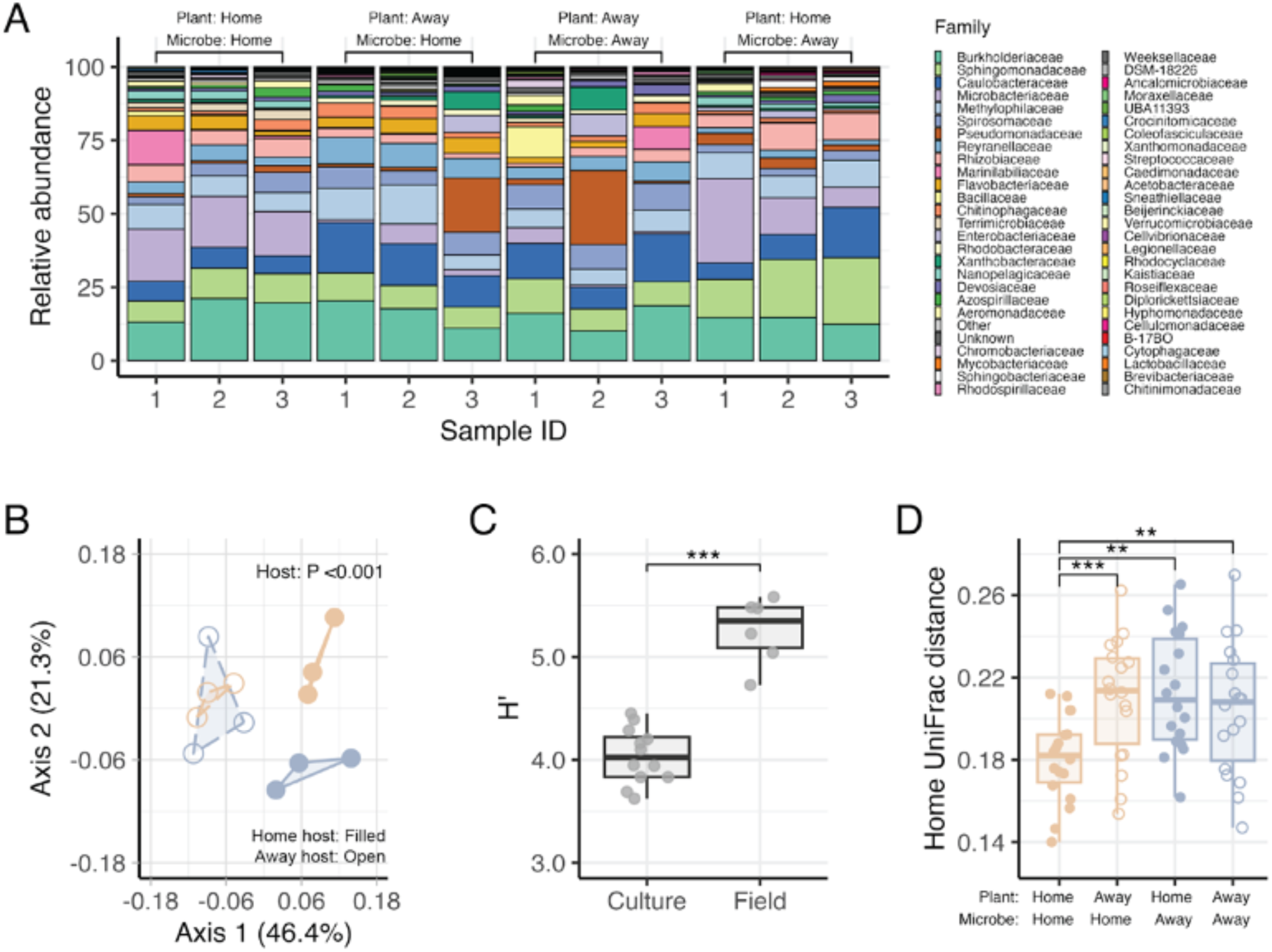
Host microbiome in source populations prior to pond introduction. (A) Family-level relative abundance of 16S rRNA gene sequences. Bars are ordered by host genotype and microbial treatment. “Other” refers to families that represent <0.1% across all samples, while “Unknown” refers to samples that could not be identified to family by sequencing. Families are ordered based on its mean abundance across all samples (B) Principal coordinates analysis (PCoA) of weighted UniFrac distances between cultured samples. Hulls are grouped by microbe treatment (home = orange; away = blue) and host genotype (home = filled circles and solid lines; away = open circles and dashed lines). (C) Estimate of Shannon index (H′) for cultured samples and field samples obtained directly from the home site. (D) Weighted UniFrac distances of cultured samples to field samples. Cultured samples are ordered by microbial treatment (home = orange; away = blue) and host genotype (home = filled circles; away = open circles). \*\*\**p*MCMC < 0.001; \*\**p*MCMC < 0.01.

Compared to lab-cultured microbiomes, the microbial diversity of field-sampled microbiomes obtained from the home site was significantly higher (mean increase in H′ = 1.172, 0.921 to 1.421, *p*MCMC < 0.001; Fig. 3C). While this suggests that some microbial diversity was lost during lab culturing, we also found similarities in taxonomic composition (Fig. S3). For example, both lab and field microbiomes supported a large proportion of Burkholderiaceae (on average representing 15.8% and 20.5% of all lab and field samples, respectively) and Sphingomonadaceae (11.2 and 5.9%; Fig. S3). Moreover, analysis of weighted UniFrac distances between lab-cultured and field microbiomes showed that home plants cultured with home microbes were most similar to field samples from the home site (Fig. 3D), with all other plant-microbe combinations having a significantly greater UniFrac distances in comparison (*p*MCMC < 0.001; Table S7).

### Host microbiomes turn over rapidly during colonization

Despite initial differences in the microbiome among source populations, we found a rapid turnover in microbiome composition once host plants were introduced into the common pond (Fig. 4A). Post-introduction, ordination analyses showed that microbiome composition changed significantly with time (PERMANOVA: F_(1,60)_ = 5.823, R^2^ = 0.080, *p* = 0.003; Table S8). We also found that from day 4 onwards, neither the initial microbial treatment (PERMANOVA: F_(2,60)_ = 0.674, R^2^ = 0.019, *p* = 0.640) nor host genotype (PERMANOVA: F_(1,60)_ = 0.588, R^2^ = 0.008, *p* = 0.604; Table S8) explained significant differences in microbiome composition. This was consistent with a rapid homogenization of the microbiome upon pond introduction, as confirmed by a weak but statistically significant decline in mean UniFrac distance (β = *–*0.002, *–*0.004 to *–*0.001, *p*MCMC = 0.007; Fig. 4B) among the microbiome of introduced hosts over time.

**FIGURE 4.**
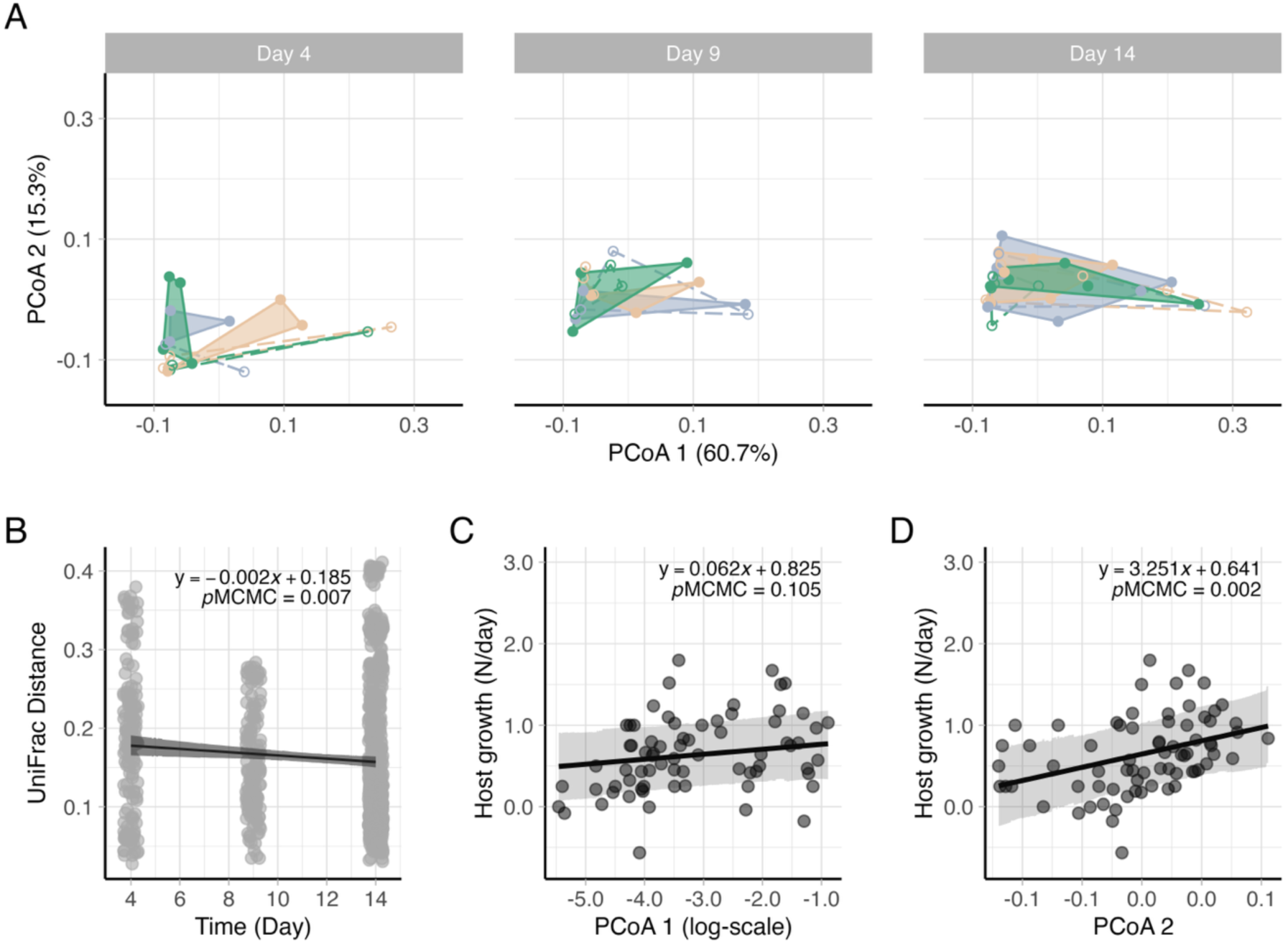
Changes in the microbiome and its effects on host performance during colonization. (A) Principal coordinates analysis (PCoA) of weighted UniFrac distances between microbial communities sampled during colonization. Hulls represent initial microbe treatment (home = orange; away = blue; green = sterile) by host genotype (home = filled circles and solid lines; away = open circles and dashed lines). The first two PCoA axes explain 76.0% of the total variation in the microbiome. (B) Change in weighted UniFrac distances between microbial communities sampled during colonization. (C; D) Relationship between PCoA axes and host performance. Data points show host growth rate (N/day) and the composition of their microbiome as represented by the first or second PCoA axis of panel A. In (C), the first PCoA axis was log-shift transformed prior to regression. Lines and bands in (B) to (D) represent model predicted mean slopes and their 95% CIs, while *p*MCMC values shown are for the slopes.

We also conducted a post-hoc ordination analysis to test if microbiome assembly on initially microbe-free hosts differed by host genotype. We found that there was a significant interaction between host genotype and time (PERMANOVA: F_(1,21)_ = 2.727, R^2^ = 0.101, *p* = 0.045; Table S9), and that microbiome composition differed by host genotype even 14 days after introduction (Fig. S4). Similarly, we conducted a post-hoc analysis to test if microbiome assembly on inoculated hosts differed by the initial microbial treatment (e.g., due to priority effects from home or away microbes), but we did not find a significant effect of initial microbial treatment (Table S10).

### Host performance in the wild is associated with microbiome composition

Along with rapid changes in the host microbiome, we found that host performance differed significantly with aspects of microbiome composition. Specifically, while host growth rates did not change significantly across the first principal coordinate analysis (PCoA) axis (β = 0.062, *–*0.013 to 0.137, *p*MCMC = 0.105; Fig. 4C), we found that host growth increased significantly with shifts in the microbiome represented by the second PCoA axis (β = 3.251, 1.231 to 5.428, *p*MCMC = 0.002; Table S11; Fig. 4D). Specifically, phylogenetic factorization identified that host performance was primarily associated with a 16-member clade of *Limnohabitans* (Burkholderiaceae family) whose relative abundance decreased with increasing host performance (*F* = 11.032; *P* = 0.001; Fig. 5; Fig. S6). In contrast, host performance was positively associated with the relative abundance of an unclassified bacterium within the Marinilabiliaceae family (*F* = 12.456; *P* < 0.001) and a clade consisting of 1451 members dominated mostly by bacteria in the orders Burkholderiales and Pseudomonadales (comprising 33.7% and 7.6% of the clade, respectively; F = 9.366; *P* = 0.003; Fig. 5).

**FIGURE 5.**
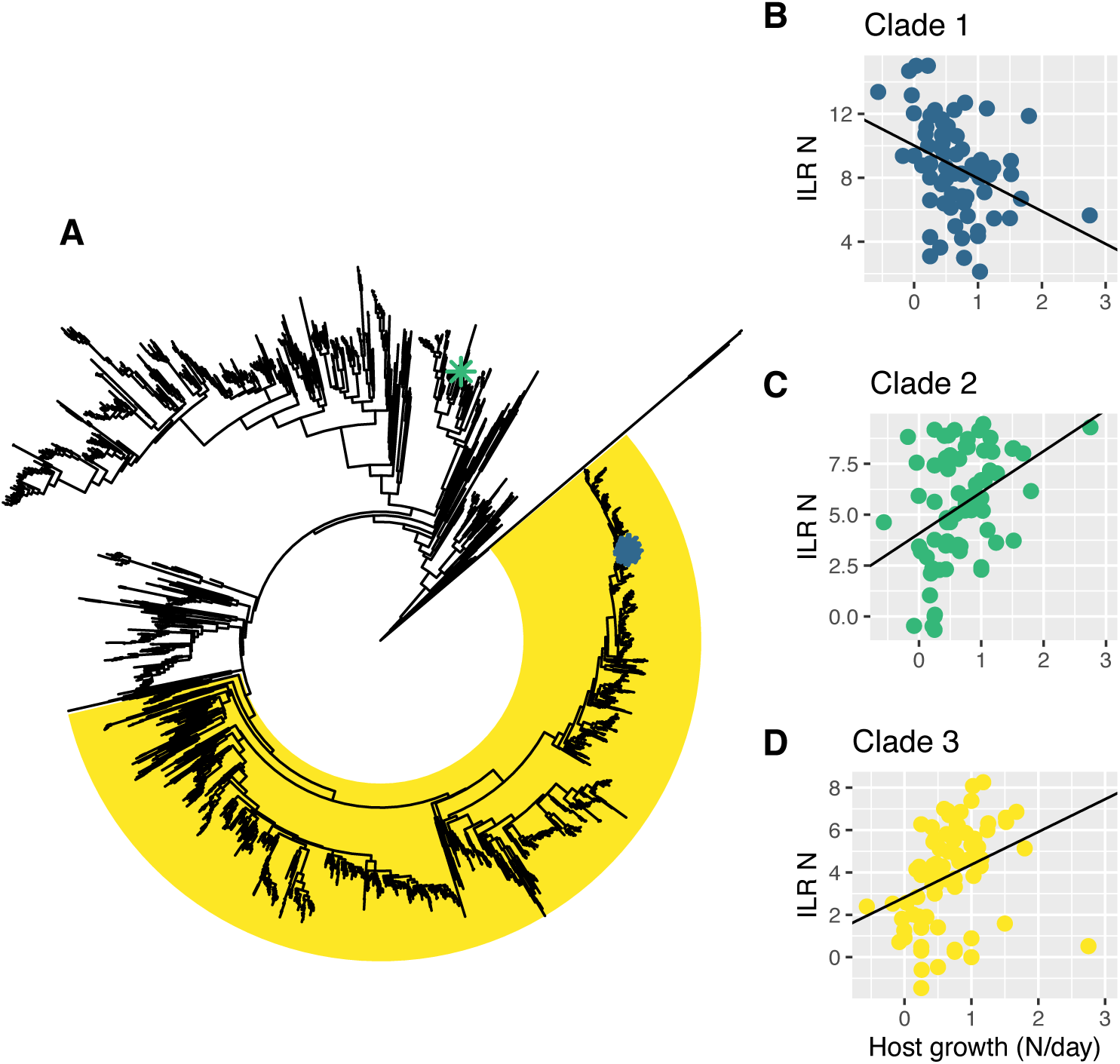
Microbial clades in the recruited microbiome that were identified to associate with host performance during colonization. (A) Phylogenetic tree of the microbiome recruited during days 4-14 of colonization in the common pond. Blue asterisks mark a 16-member clade of bacteria in the *Limnohabitans* genus (Burkholderiaceae family) while the green asterisk marks an unidentified taxon in the Marinilabiliaceae family. The yellow highlight is a clade of 1451 members dominated mostly by bacteria in the orders Burkholderiales and Pseudomonadales. (B) to (D) show the linear relationship between the isometric log-ratio (ILR) transformed mean abundance (*y*-axis) of each clade identified in (A) to host performance (*x*-axis) as obtained by phylogenetic factorization.

## DISCUSSION

Understanding the processes that alter colonization success is fundamental to predicting future biological introductions and changes to species distributions (Angert et al. 2020; Iseli et al. 2023). With increasing evidence suggesting that host-microbe mutualisms play a critical role in shaping species distributions (Delevaux et al. 2024; Nathan et al. 2023), we set out to test empirically whether and how microbiomes altered host performance during experimental co-introductions in the wild. Overall, our results demonstrate that while microbes recruited during colonization can influence host performance, the identity of the initial microbiome that plants are co-introduced with can also significantly alter the trajectory of host growth. Co-introduced symbionts can therefore leave lasting impacts on host performance even if the microbiome rapidly turns over during colonization.

Notably, hosts that were co-introduced with home microbes (i.e., local to the habitat being colonized) grew faster compared to hosts co-introduced with away (i.e., non-local) microbes, with effects consistent across host genotype (Fig. 2). This suggests that home microbes are matched to abiotic or biotic conditions of their home site (Johnson et al. 2010; Rúa et al. 2016) and can provide greater benefits to hosts than non-local microbes (i.e., a G_microbe_ x E interaction), or that microbes from the home site are generally better at provisioning host benefits than away microbes in our experiment (i.e., a main effect of G_microbe_). Additionally, we also found that when hosts were introduced without a microbiome, home host genotypes grew faster than away genotypes, suggesting that local hosts are also matched to conditions at their home site (O’Brien et al. 2024), or that local hosts introduced without microbes quickly acquire microbial partners that are different to non-local plants (Fig. S4) (Wippel et al. 2021). In the context of colonization, these results suggest that host performance and establishment may be hindered by mismatches between the introduced environment and the genetic background of both hosts and co-introduced microbes.

Greater host performance with home microbes also suggests that introduced host populations could colonize faster if they are able to rapidly acquire local microbial symbionts after introduction. While founder populations harboured distinct microbiomes prior to introduction (Fig. 3), we found that these initial differences were rapidly homogenized upon introduction (Fig. 4A and 4B). We therefore did not find that microbes that were initially present on hosts prevented or altered the assembly of later-arriving microbes through priority effects or competitive exclusion, as previously reported in other symbiotic microbes (Boyle et al. 2021; Chappell et al. 2022). Instead, this rapid turnover in the microbiome is consistent with the prediction that ‘ecological fitting’ (i.e., the acquisition of local microbes during colonization) is common for hosts that have diffuse and generalized interactions with their microbes (Le Roux et al. 2017) such as found in duckweeds (Laurich et al. 2024). Consistently, we found significant associations between host performance and the composition of the recruited microbiome during colonization (Fig. 4C-D), and that host performance was associated with specific bacterial clades that have been commonly found in the duckweed microbiome and in freshwater ecosystems (Fig. 5) (Anneberg et al. 2023; Baggs et al. 2022; Kasalický et al. 2013). We also found similar associations between host performance and the same (or closely related) clades across hosts with different initial microbiomes (Fig. S7). This suggests that once hosts rapidly recruited microbes from the introduced environment, similar subsets of microbial taxa affected host growth regardless of the host’s initial microbiome.

Despite this rapid homogenization of the microbiome during colonization, we found that hosts differed significantly in their performance, and that the initial microbiome that hosts were co-introduced with had persistent effects on host growth throughout the experiment (Fig. 2). Furthermore, while host performance was more variable among plants when their microbiomes were less similar to one another, we found that this relationship was strongest early in the colonization process and faded over time (Fig. S5). These results suggest that the initial microbiome left a persistent, physiological imprint on host plants that lasted for multiple host generations (4-5 generations assuming ∼3 day generation time) (Ziegler et al. 2015). Transgenerational responses to stress have previously been shown to last for up to 12 generations in duckweeds (Van Antro et al. 2023). Microbially mediated transgenerational effects have also been found to improve performance in plants associating with mycorrhizal fungi (Latzel et al. 2025; Puy et al. 2022). Such effects suggest that non-local microbes that are co-introduced with hosts can leave a lasting impact on host population dynamics, even in the case of generalized mutualisms where hosts can subsequently recruit local microbes during colonization.xs

Overall, our results emphasize that the host colonization success could be strongly influenced by the initial host microbiome, regardless of how fast hosts acquire new microbial partners in the introduced habitat. Future studies should use a reciprocal transplant design to tease apart the effects of host and microbe genotype by environment interactions, and test co-introductions across larger spatial and environmental scales (e.g., beyond the current range) (Benning & Moeller 2021). Nonetheless, our current study suggests that understanding the spread of invasive plants or the success of transplanted agricultural crops should consider that microbiome co-introduction could significantly influence the outcome of establishing populations. For duckweeds, this is particularly relevant given that some species are rapidly invading (Andrade-Pereira 2024), while other species are being introduced into freshwater habitats for applications in remediation (Zhou et al. 2023). In conclusion, understanding the intimate association between plant and microbes are crucial for predicting the rate of colonization in this rapidly changing world.

## ACKNOWLEDGEMENTS

We thank members of the Frederickson Lab for feedback, and thank C.I. Carlson, R. Molnarova, O. Pogoutse, and K.D. Ricks for lab and field assistance. We thank T. Zallek for guidance on how to construct the floating platforms. This work was funded by a Koffler Scientific Reserve Undergraduate Student Research Award (KSR USRA) to J.Y. Funding from the Faculty of Arts & Sciences Postdoctoral Fellowship supported T.U. Funding from a Natural Sciences and Engineering Research Council of Canada (NSERC) Discovery Grant, the Gordon & Betty Moore Foundation (Grants GBMF9536 [doi.org/10.37807/GBMF9356] and GBM10635 [doi.org/10.37807/GBMF10635]), and the National Science Foundation (Award number 2300057) supported M.E.F.

## Statement of authorship

T.U. and M.E.F. designed the study in collaboration with J.Y. J.Y. and T.U. conducted the experiment. J.Y. sequenced the microbes. J.Y. led the initial analysis and writing, with feedback from T.U. and M.E.F. T.U. revised the manuscript with feedback from J.Y. and M.E.F. All authors contributed to final revisions and editing.

## Notes

### Competing Interest Statement

The authors have declared no competing interest.

