## Supplementary Information for "For colonization success, should hosts and microbes travel alone, together, or swap partners along the way?"

Supplementary Figures

Figure S1. Sensitivity analysis of host performance
Figure S2. Phylogenetic tree of the initial host microbiome
Figure S3. Family-level bar plots of 16s rRNA sequences (lab and field samples)
Figure S4. PCoA of the microbiome during colonization for initially sterile hosts
Figure S5. Relationship between host growth and microbiome composition over time
Figure S6. Ordination of ILR coordinates for clades in the recruited microbiome
Figure S7. Association of clades in the recruited microbiome with host performance

Supplementary Tables

Table S1. V3-V4 region of the 16s rRNA gene
Table S2-S5. Model coefficients for analysis of host performance
Table S6. PERMANOVA of initial microbiome differences
Table S7. Model coefficients for comparison of lab and field microbiome samples
Table S8. PERMANOVA of microbiome during colonization
Table S9. PERMANOVA of microbiome during colonization (initially sterile hosts)
Table S10. PERMANOVA of microbiome during colonization (inoculated hosts)
Table S11. Model coefficients for analysis of host growth by microbiome composition

### Supplementary Tables

**Table S1:** Forward and reverse primers used in the amplification of the V3-V4 region in the 16S rRNA gene.

| Primer | Sequence |
|---|---|
| Forward (341F) | CCTACGGGNGGCWGCAG<br>TCCTACGGGNGGCWGCAG<br>ACCCTACGGGNGGCWGCAG<br>CAACCTACGGGNGGCWGCAG |
| Reverse (805R) | GACTACHVGGGTATCTAATCC<br>TGACTACHVGGGTATCTAATCC<br>ACGACTACHVGGGTATCTAATCC<br>CATGACTACHVGGGTATCTAATCC |

**Table S2:** Model coefficients for host population growth rate. Posterior represents the mean estimate, while LCI and UCI represent the lower and upper credible intervals, respectively. Random effects included pair ID, platform ID, and replicate ID. *Italicized* estimates have 95% credible intervals that do not span zero. Intercept represents the number of individuals on Day 0 for ‘home’ host plants inoculated with ‘home’ microbes.

| Fixed effect predictors | Posterior | LCI | UCI | *p*MCMC |
|---|---:|---:|---:|---:|
| *Intercept* | *9.170* | *6.931* | *11.483* | *<0.001* |
| *Day* | *0.781* | *0.710* | *0.851* | *<0.001* |
| Host (away) | –0.241 | –1.326 | 0.824 | 0.646 |
| Microbe (away) | 0.396 | –0.656 | 1.473 | 0.467 |
| Microbe (none) | 0.471 | –0.561 | 1.583 | 0.393 |
| Day x Host (away) | 0.027 | –0.071 | 0.129 | 0.598 |
| *Day x Microbe (away)* | *–0.144* | *–0.238* | *–0.044* | *0.002* |
| Day x Microbe (none) | 0.026 | –0.070 | 0.128 | 0.598 |
| Host (away) x Microbe (away) | 0.260 | –1.218 | 1.788 | 0.737 |
| Host (away) x Microbe (none) | 0.159 | –1.316 | 1.709 | 0.838 |
| Day x Host (away) x Microbe (away) | –0.035 | –0.167 | 0.103 | 0.616 |
| *Day x Host (away) x Microbe (none)* | *–0.289* | *–0.430* | *–0.150* | *<0.001* |

**Table S3:** Contrast model coefficients for host population growth rate. Posterior represents the mean estimate, while LCI and UCI represent the lower and upper credible intervals, respectively. Random effects included pair ID, platform ID, and replicate ID. *Italicized* estimates have 95% credible intervals that do not span zero. Intercept represents the number of individuals on Day 0 for 'away' host plants inoculated with 'home' microbes.

| Fixed effect predictors | Posterior | LCI | UCI | *p*MCMC |
|---|---:|---:|---:|---:|
| *Intercept* | *8.924* | *6.806* | *11.259* | *<0.001* |
| *Day* | *0.807* | *0.737* | *0.876* | *<0.001* |
| Host (home) | 0.243 | –0.865 | 1.265 | 0.661 |
| Microbe (away) | 0.647 | –0.388 | 1.700 | 0.232 |
| Microbe (none) | 0.616 | –0.436 | 1.664 | 0.256 |
| Day x Host (home) | –0.027 | –0.124 | 0.073 | 0.591 |
| *Day x Microbe (away)* | *–0.178* | *–0.273* | *–0.081* | *<0.001* |
| *Day x Microbe (none)* | *–0.261* | *–0.354* | *–0.160* | *<0.001* |
| Host (home) x Microbe (away) | –0.253 | –1.668 | 1.263 | 0.731 |
| Host (home) x Microbe (none) | –0.164 | –1.672 | 1.361 | 0.821 |
| Day x Host (home) x Microbe (away) | 0.033 | –0.101 | 0.171 | 0.642 |
| *Day x Host (home) x Microbe (none)* | *0.288* | *0.154* | *0.435* | *<0.001* |

**Table S4:** Contrast model coefficients for host population growth rate. Posterior represents the mean estimate, while LCI and UCI represent the lower and upper credible intervals, respectively. Random effects included pair ID, platform ID, and replicate ID. *Italicized* estimates have 95% credible intervals that do not span zero. Intercept represents the number of individuals on Day 0 for 'home' host plants that were initially uninoculated.

| Fixed effect predictors | Posterior | LCI | UCI | *p*MCMC |
|---|---:|---:|---:|---:|
| *Intercept* | *9.674* | *7.349* | *11.924* | *<0.001* |
| *Day* | *0.806* | *0.736* | *0.873* | *<0.001* |
| Host (away) | –0.102 | –1.145 | 0.993 | 0.862 |
| Microbe (away) | –0.089 | –1.098 | 1.025 | 0.869 |
| Microbe (home) | –0.475 | –1.543 | 0.594 | 0.381 |
| *Day x Host (away)* | *–0.260* | *–0.356* | *–0.161* | *<0.001* |
| *Day x Microbe (away)* | *–0.169* | *–0.264* | *–0.068* | *0.001* |
| Day x Microbe (home) | –0.025 | –0.123 | 0.073 | 0.615 |
| Host (away away) x Microbe (away) | 0.119 | –1.332 | 1.593 | 0.882 |
| Host (home) x Microbe (home) | –0.147 | –1.583 | 1.454 | 0.836 |
| *Day x Host (away) x Microbe (away)* | *0.253* | *0.111* | *0.390* | *<0.001* |
| *Day x Host (away) x Microbe (home)* | *0.288* | *0.154* | *0.434* | *<0.001* |

**Table S5:** Sensitivity model coefficients for host population growth rate (including plant populations destructively harvested during the experiment for micorbiome data). Posterior represents the mean estimate, while LCI and UCI represent the lower and upper credible intervals, respectively. Random effects included pair ID, platform ID, and replicate ID. *Italicized* estimates have 95% credible intervals that do not span zero. Intercept represents the number of individuals on Day 0 for 'home' host plants inoculated with 'home' microbes.

| Fixed effect predictors | Posterior | LCI | UCI | *p*MCMC |
|---|---|---|---|---|
| *Intercept* | *9.194* | *7.305* | *11.055* | *<0.001* |
| *Day* | *0.794* | *0.729* | *0.858* | *<0.001* |
| Host (away) | –0.060 | –1.010 | 0.821 | 0.906 |
| Microbe (away) | 0.422 | –0.435 | 1.394 | 0.354 |
| Microbe (none) | 0.414 | –0.483 | 1.371 | 0.376 |
| Day x Host (away) | –0.004 | –0.097 | 0.087 | 0.943 |
| *Day x Microbe (away)* | *–0.155* | *–0.246* | *–0.066* | *0.001* |
| Day x Microbe (none) | 0.010 | –0.085 | 0.105 | 0.834 |
| Host (away) x Microbe (away) | 0.255 | –1.008 | 1.540 | 0.708 |
| Host (away) x Microbe (none) | 0.024 | –1.360 | 1.244 | 0.968 |
| Day x Host (away) x Microbe (away) | –0.018 | –0.151 | 0.106 | 0.775 |
| *Day x Host (away) x Microbe (none)* | *–0.246* | *–0.373* | *–0.113* | *<0.001* |

**Table S6:** PERMANOVA output for host microbiome composition prior to introduction. Italicised estimates have $P < 0.05$. DF = degrees of freedom; SS = sum of squares.

| | DF | SS | $R^2$ | F | *P* |
|---|---|---|---|---|---|
| *Host* | *1* | *0.077* | *0.402* | *7.904* | *<0.001* |
| Microbe | 1 | 0.019 | 0.098 | 1.928 | 0.107 |
| Host x Microbe | 1 | 0.018 | 0.093 | 1.831 | 0.121 |
| Residual | 8 | 0.078 | 0.407 | | |
| Total | 11 | 0.193 | 1.000 | | |

**Table S7:** Model coefficients of weighted UniFrac distances measuring similarity of the lab-cultured host microbiomes to that of microbiomes from field plants of the home site. Posterior represents the mean estimate, while LCI and UCI represent the lsower and upper credible intervals, respectively. *Italicized* estimates have 95% credible intervals that do not span zero. Intercept represents the weighted UniFrac distance of 'home' host plants and microbes.

| Fixed effect predictors | Posterior Mean | LCI | UCI | *p*MCMC |
|---|---|---|---|---|
| *Intercept* | *0.180* | *0.167* | *0.194* | *<0.001* |
| *Microbe (Away)* | *0.032* | *0.013* | *0.051* | *<0.001* |
| Host (Away) | 0.029 | 0.010 | 0.048 | 0.004 |
| Host x Microbe | –0.037 | –0.063 | –0.010 | 0.007 |

**Table S8:** PERMANOVA output of microbiome composition during colonization across all samples. Italicised estimates have $P < 0.05$. DF = degrees of freedom; SS = sum of squares.

| | Df | SS | $R^2$ | F | *P* |
|---|---|---|---|---|---|
| *Day* | *1* | *0.108* | *0.080* | *5.823* | *0.003* |
| Microbe | 2 | 0.025 | 0.019 | 0.674 | 0.640 |
| Host | 1 | 0.011 | 0.008 | 0.588 | 0.604 |
| Day x Microbe | 2 | 0.008 | 0.006 | 0.229 | 0.986 |
| Day x Host | 1 | 0.018 | 0.013 | 0.955 | 0.374 |
| Microbe x Host | 2 | 0.012 | 0.009 | 0.324 | 0.947 |
| Day x Microbe x Host | 2 | 0.050 | 0.037 | 1.337 | 0.230 |
| Residual | 60 | 1.111 | 0.828 | | |
| Total | 71 | 1.342 | 1.000 | | |

**Table S9:** PERMANOVA output of microbiome composition during colonization for initially sterile host plants only. Italicised estimates have $P < 0.05$. DF = degrees of freedom; SS = sum of squares.

| | Df | SS | $R^2$ | F | *P* |
|---|---|---|---|---|---|
| Day | 1 | 0.036 | 0.095 | 2.547 | 0.055 |
| Host | 1 | 0.009 | 0.024 | 0.634 | 0.612 |
| *Day x Host* | *1* | *0.039* | *0.101* | *2.727* | *0.045* |
| Residual | 21 | 0.297 | 0.780 | | |
| Total | 24 | 0.380 | 1.000 | | |

**Table S10:** PERMANOVA output of microbiome composition during colonization for host plants inoculated with CDV or KSR microbes only. Italicised estimates have $P < 0.05$. DF = degrees of freedom; SS = sum of squares.

| | Df | SS | $R^2$ | F | *P* |
|---|---|---|---|---|---|
| *Day* | *1* | *0.072* | *0.077* | *3.623* | *0.026* |
| Microbe | 1 | 0.006 | 0.006 | 0.294 | 0.867 |
| Day x Microbe | 1 | 0.008 | 0.009 | 0.403 | 0.758 |
| Residual | 43 | 0.857 | 0.909 | | |
| Total | 46 | 0.943 | 1.000 | | |

**Table S11:** Model coefficients for change in host growth by the first two PCoA axes. Each PCoA axis was fit as a separate model. Posterior represents the mean estimate, while LCI and UCI represent the lower and upper credible intervals, respectively. *Italicized* estimates have 95% credible intervals that do not span zero.

| Fixed effect predictors | Posterior Mean | LCI | UCI | *p*MCMC |
|---|---|---|---|---|
| *Intercept* | *0.825* | *0.378* | *1.251* | *0.005* |
| PCoA1 | 0.062 | –0.013 | 0.137 | 0.105 |
| *Intercept* | *0.641* | *0.270* | *1.049* | *0.010* |
| *PCoA2* | *3.251* | *1.231* | *5.428* | *0.002* |

### Supplementary Figures

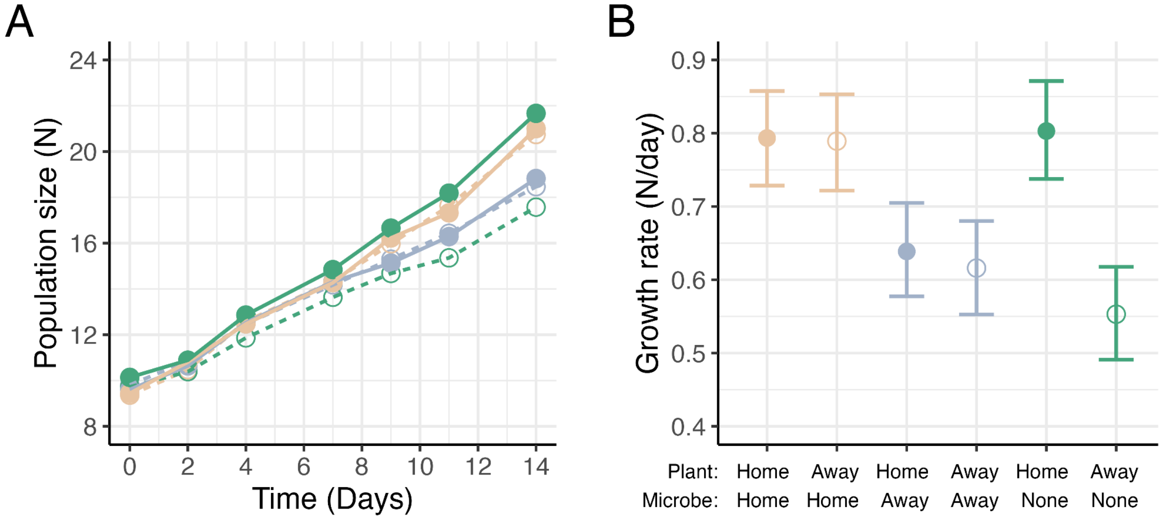

**Figure S1:** Sensitivity analysis of host growth rates, including data from plants that were destructively harvested. Home plants = filled circles and solid lines; Away plants = open circles and dashed lines; Home microbes = orange; Away microbes = blue; Uninoculated = green. (A) Lines represent the growth trajectory of introduced host populations over time where circles represent the mean population size at a given day for each plant-microbe combination. (B) Mean population growth rate (N/day) and 95% CIs for plant-microbe combination, as estimated from linear mixed-effects models (see Table S5 for model coefficients).

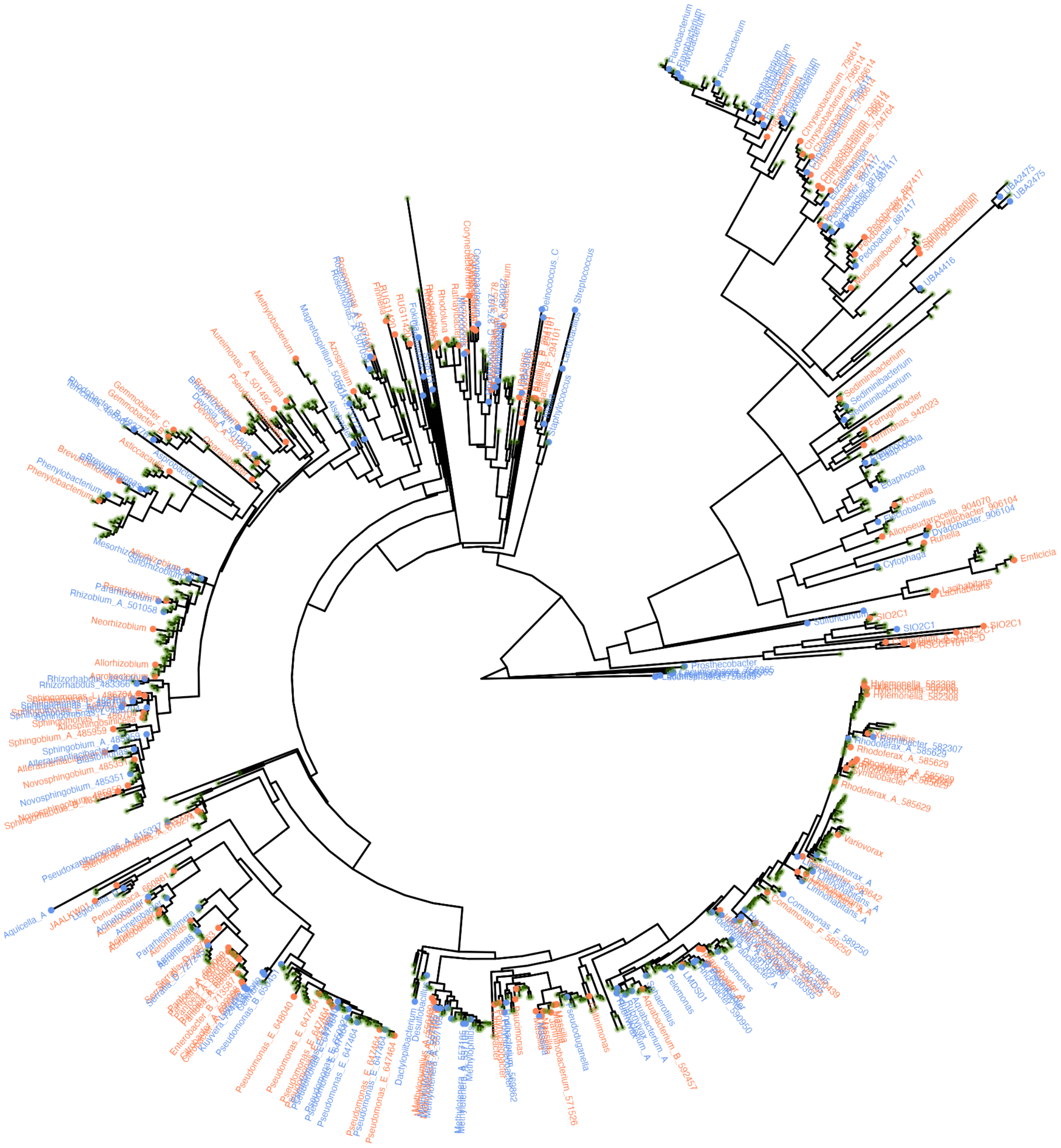

**Figure S2:** Phylogenetic tree of the initial plant microbiome before pond introduction. Green circles represent ASVs that were present in both KSR ('home') and CDV ('away') microbial treatments, while blue and orange circles represent ASVs that were distinct to either KSR or CDV microbes, respectively. For clarity, genus names are shown only for ASVs that were distinct to either KSR or CDV microbes, in blue and orange text, respectively.

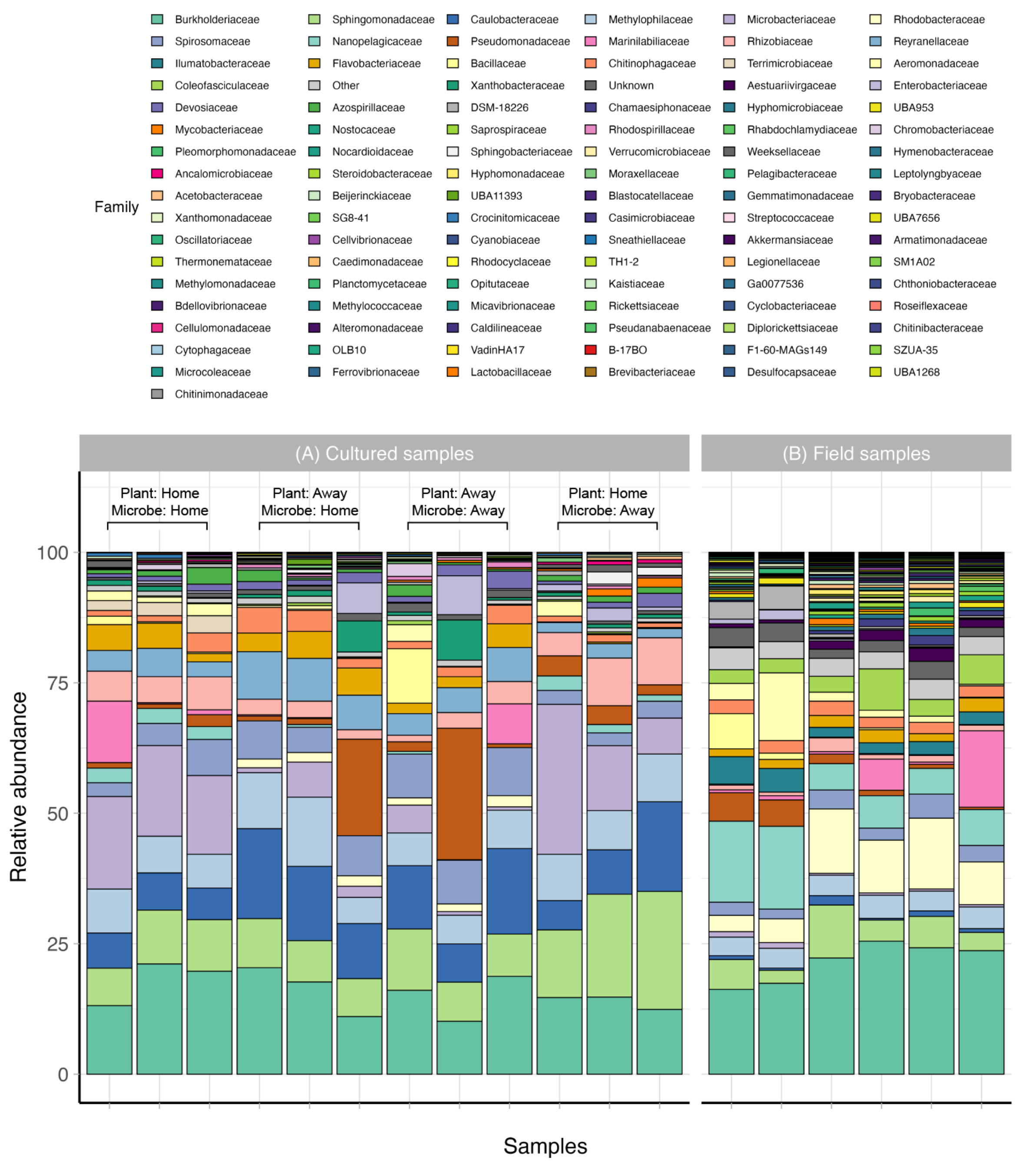

**Figure S3:** Family-level, relative abundance of 16S rRNA gene sequences obtained from (A) host microbiomes after 14 days of lab culturing prior to introduction; and (B) host microbiome samples obtained from the field (at the home site) on day 0. Bars are ordered by host genotype and microbial treatment. “Other” refers to families that represent <0.1% across all samples, while “Unknown” refers to samples that could not be identified by sequencing to family level.

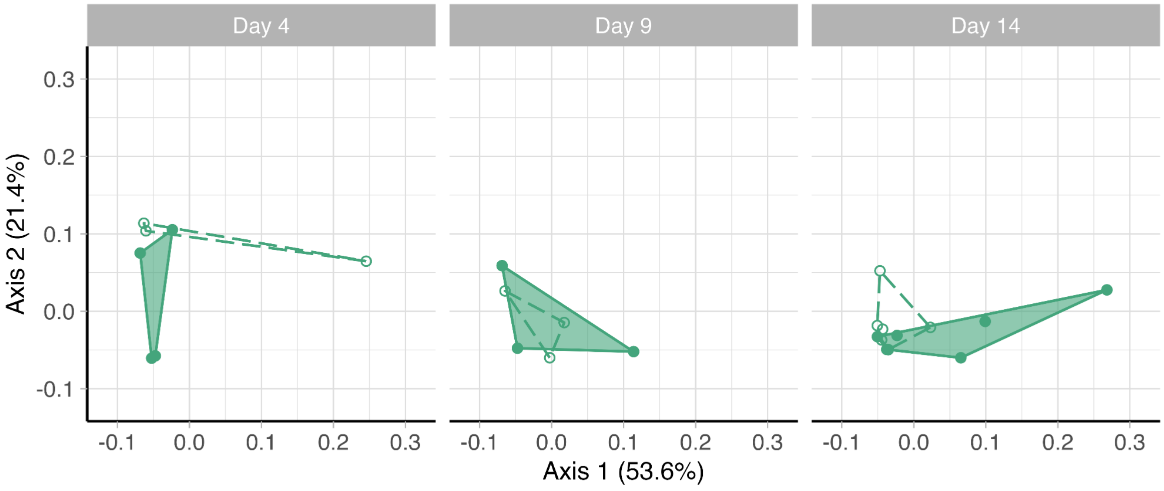

**Figure S4.** Principal coordinates analysis (PCoA) of microbiome composition (weighted UniFrac distance) during colonization for initially sterile plants only. Hulls are grouped by host genotype (home = filled circles and solid lines; away = open circles and dashed lines). See Table S10 for model coefficients.

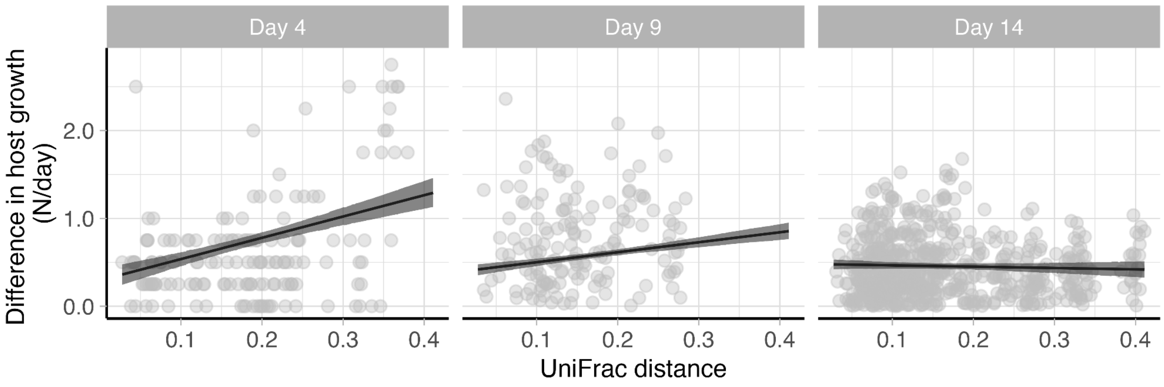

**Figure S5.** Relationship between pairwise differences in host growth rate (N/day) and host microbiome composition (weighted UniFrac distance) over time. Host plants that harbour increasingly different microbiomes have larger differences in host growth rate on day 4 ($\beta$ = 2.424, 95% CI: 1.764 to 3.112), yet there is a significant interaction between UniFrac distance and time (*p*MCMC < 0.001) such that this relationship weakens over time. By day 14, there is no longer a positive relationship between pairwise differences in host growth and UniFrac distance ($\beta$ = –0.153, 95% CI: –0.474 to 0.210).

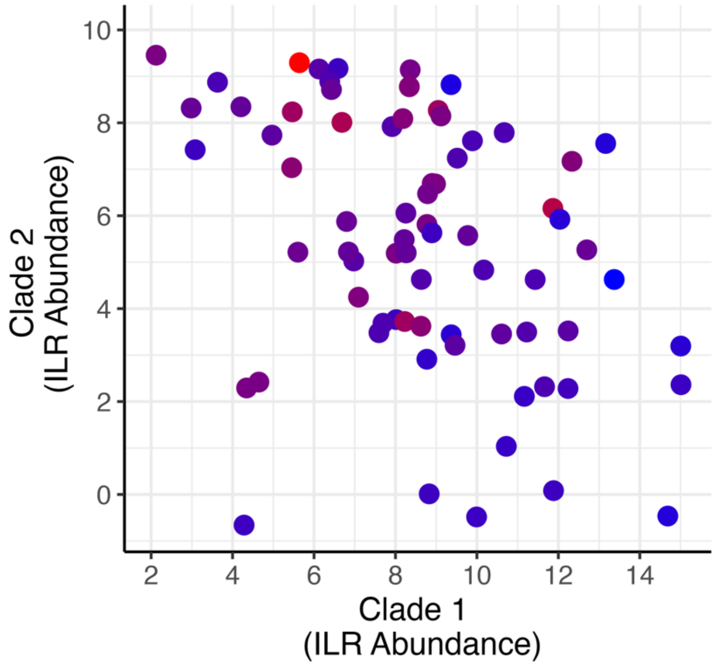

**Figure S6.** Ordination of isometric log-ratio (ILR) coordinates for clades 1 and 2 as identified by phylogenetic factorization of the microbiome recruited during colonization. The colour gradient represents host performance, where blue and red represent lower and higher host growth rates, respectively.

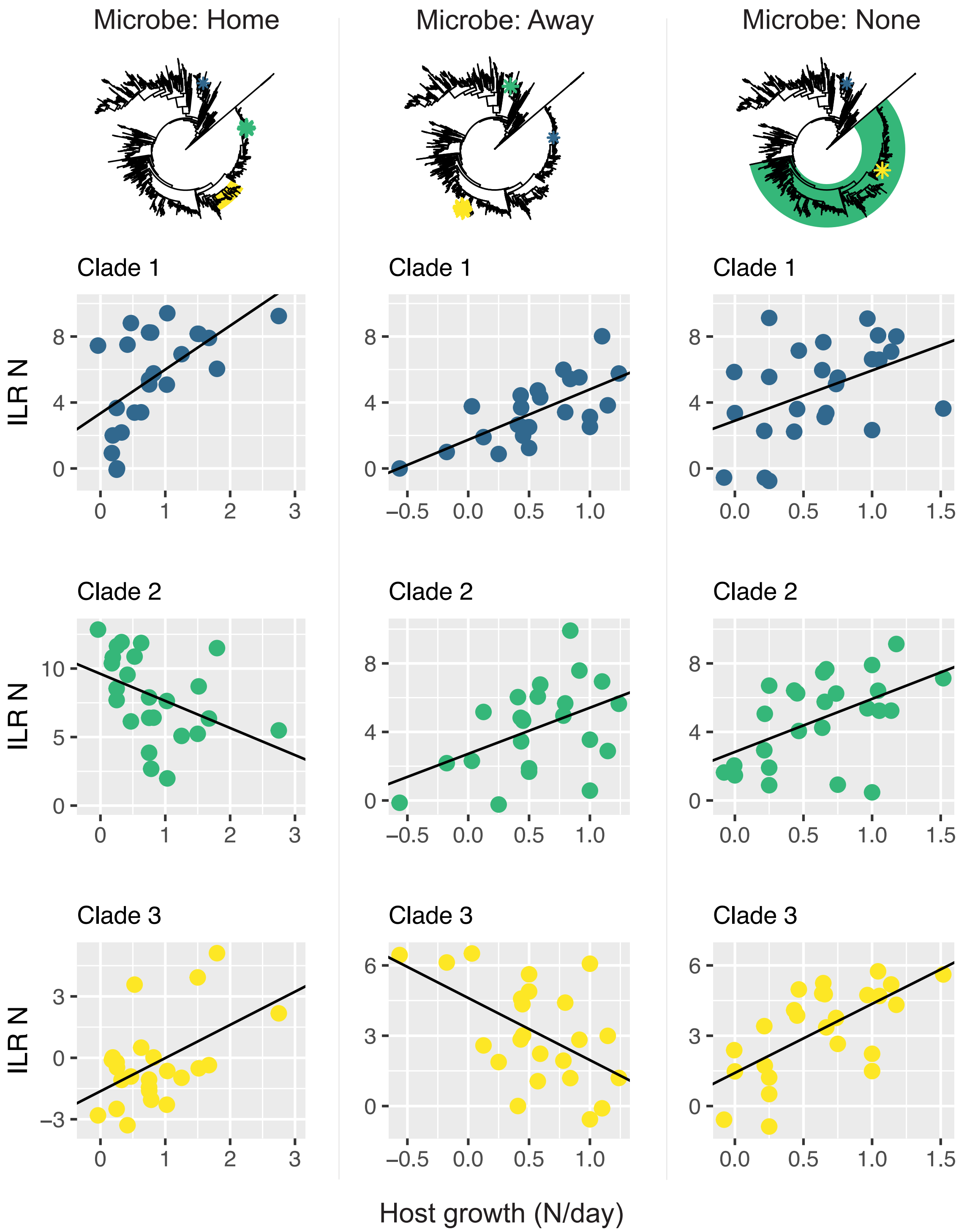

**Figure S7.** Clades in the recruited microbiome that were identified to associate with host performance during colonization. Clades (Clade 1 = blue; Clade 2 = green; Clade 3 = yellow) that associate with host performance have been separately identified for hosts that initially differed in their co-introduced microbiome (column 1 = hosts initially co-introduced with ‘home’ microbiome; column 2 = hosts initially co-introduced with ‘away’ microbiome; column 3 = hosts initially introduced without a microbiome). Note in the tree that the same or phylogenetically similar clades are identified to be important for host growth across all hosts. Panels show the linear relationship between the isometric log-ratio (ILR) transformed mean abundance (*y*-axis) of each clade to host performance (*x*-axis) as obtained by phylogenetic factorization. Note that the *y*-axes and *x*-axes are scaled differently across panels.
